# Scalable neighbour search and alignment with uvaia

**DOI:** 10.1101/2023.01.31.526458

**Authors:** Leonardo de Oliveira Martins, Alison E. Mather, Andrew J. Page

## Abstract

Despite millions of SARS-CoV-2 genomes being sequenced and shared globally, manipulating such data sets is still challenging, especially selecting sequences for focused phylogenetic analysis. We present a novel method, uvaia, which is based on partial and exact sequence similarity for quickly extracting database sequences similar to query sequences of interest. Many SARS-CoV-2 phylogenetic analyses rely on very low numbers of ambiguous sites as a measure of quality since ambiguous sites do not contribute to single nucleotide polymorphism (SNP) differences, which uvaia alleviates by using measures of sequence similarity that consider partially ambiguous sites. Such fine-grained definition of similarity allows not only for better phylogenetic analyses, but also for improved classification and biogeographical inferences. Uvaia works natively with compressed files, can use multiple cores and efficiently utilises memory, being able to analyse large data sets on a standard desktop.

## Introduction

Genome sequencing has been globally deployed at pace to understand the evolution, transmission and dynamics of the SARS-CoV-2 virus, with the goal of providing actionable data for management of the COVID-19 pandemic (du Plessis *et al*., 2021; Maxmen, 2021; Lambrou *et al*., 2022). Genomic epidemiology places genomes into context, with the most basic questions being, is a genome similar or different to what has been seen before? The next is, what are the most similar genomes? This can allow for outbreaks to be identified, linked, and mitigations put in place to monitor or limit further spread. This is particularly important for closed environments such as hospitals (Page *et al*., 2021), care homes (Aggarwal *et al*., 2021), or for limiting the spread of newly emergent variants with concerning mutations (Aggarwal *et al*., 2022).

The number of SARS-CoV-2 genomes available in global public databases such as the ENA/NCBI and GISAID has surpassed 14 million (accessed 2023-01-23). SARS-CoV-2 is now the most sequenced organism of all time, however bioinformatics methods have struggled to keep pace with the scale of the data, or are optimised for different properties (e.g. small numbers of large genomes, rather than large numbers of small genomes). To complicate things further, the sequencing methods commonly utilised for SARS-CoV-2 can result in partial genomes (Baker *et al*., 2021). This can be due to a low viral load of the sample where a patient is at an early or late stage of their infection, due to a mutation causing a dropout in an amplicon primer sequence (Sanderson and Barrett, 2021), or could be due to the manner the sample has been collected and stored (Liu *et al*., 2021).

Another algorithmic challenge is that SARS-CoV-2 mutates relatively slowly, with regular global lineage replacements as fitter lineages emerge, so circulating diversity can be very low during any given time period, making every mutation count. To address huge genomic SARS-CoV-2 data sets with uneven quality, resolution, and completeness, uvaia was created so that it only uses the sequence data and similarity measures ―which indirectly account for genome completeness, as we’ll see below.

To account for partial genomes, a minimum threshold for genome completeness is often applied, so that algorithms only need to focus on high quality genomes. For example, the COVID-19 Genomics UK (COG-UK) consortium sets a threshold of 50% (Page *et al*., 2021), with data deposited in the ENA/NCBI and GISAID (>90% completeness). To maximise the chances of getting a high coverage genome, often a cycle threshold (Ct) is applied even before sequencing begins: the Ct value is the number of polymerase chain reaction amplification cycles necessary for the virus to be detected (Rhoads *et al*., 2021). Thus higher Ct values mean that there is less viral RNA, and samples with Ct > 30 are usually excluded, to ensure that there is sufficient viral material available for sequencing. Phylogenetic analyses can decide for a higher genome completeness threshold, e.g. at least 90% of the sites with high-coverage. This can result in clinically important samples being disregarded before sequencing, and for sequenced genomes to never be made public, even though they could contain epidemiologically useful information. Assuming that the sequence completeness does not compromise the alignment (i.e. the presence of ambiguous sites does not generate a spurious alignment with regards to the reference genome), uvaia can find similar sequences based on the number of partial and exact matches, instead of the more common number of single nucleotide polymorphism (SNP) differences.

Two important software tools for the phylogenetic analysis of SARS-CoV-2 data sets are Nextrain (Hadfield *et al*., 2018) and civet (O’Toole *et al*., 2021), and both contain strategies for reducing the number of sequences to those relevant to a particular study. Nextstrain can generate “focal sets” by downsampling based on geography and collection date information, which can subsequently be selected for inclusion through a genetic distance-based “priority” ordering (Hodcroft *et al*., 2021). Likewise, given a background database composed of alignment and metadata, civet can generate a “catchment area” composed of sequences within a given distance to the query genomes, with the possibility of downsampling (O’Toole *et al*., 2021). Both methods rely on stochastic factors, metadata information and a SNP-distance measure.

Another essential software for SARS-CoV-2 analysis with integrated phylogenetics is UShER, which can parsimoniously place a query sequence into a tree, and therefore can return the closest samples from this so-called mutation-annotated tree (Turakhia *et al*., 2021). It neglects indels, depends on sequences in the VCF format, and relies on the existence of the maximum parsimony tree. Nonetheless it has good accuracy for PANGO lineage classification and scales very well (Thornlow *et al*., 2021).

We have created a method, uvaia, for performing neighbourhood search and alignment, allowing for similar genomes to be found within massive datasets. Given a genome, we can rapidly find all other similar, equivalent or identical genomes from massive public datasets. Uvaia scales linearly to the massive datasets seen with SARS-CoV-2 genomics, and linearly per core to the query sample size. It accounts for the complexity of partial genomes, and can provide analysis rapidly on an ordinary laptop.

Uvaia has been used for rapidly analysing genomic data to assist pandemic management in multiple countries, including distance based analyses analysing dynamics of the rapid emergence of the Omicron variant of concern in the UK (Eales, de Oliveira Martins, *et al*., 2022), for phylogenetic based analyses to understand multiple waves in Zimbabwe (Mashe, Takawira, Gumbo, *et al*., 2021), the spread of variants of concern in Pakistan (Sarwar *et al*., 2021), the emergence and replacement of multiple variants of concern in Lebanon (Merhi *et al*., 2022). Uvaia achieves this by utilising high efficiency compression, efficient parallelisation and combines this with knowledge of the fundamental characteristics and properties found with SARS-CoV-2 genomic datasets. Uvaia is available under the open source GNU GPL3 licence from https://github.com/quadram-institute-bioscience/uvaia.

Uvaia tries to solve two problems related to massive data sets, which have not been fully explored by existing tools: by working natively with XZ-compressed files, and by working with a pool of sequences in parallel. The first is to avoid files which may compromise the disk space: the raw GISAID fasta file with all sequences as of v.2022_04_11, with more than 10Mi sequences, occupies 282.2 GiB, while its XZ-compressed equivalent takes only 919.2 MiB. Such files cannot be read into memory at once in most personal computers either, and thus the uvaia programs work with the sequence files in manageable batches, using multiprocessing whenever possible, including the output XZ compression which is done in parallel.

## Handling ambiguity

Most SARS-CoV-2 sequences will have some indels but also a considerable number of Ns, which represent complete uncertainty or ambiguity in the base at the location. A sequence may also have partially ambiguous sites (IUPAC codes), as for instance the character M means that the site may be an A or a C, but not a G or a T. Both gaps and Ns are excluded from pairwise comparison by most sequence comparison algorithms, including uvaia. Uvaia calculates the pairwise similarity based on unambiguous pairwise comparisons (i.e. sites exclusively with A, C, G, or T on both sequences), on partial matches (so that an M will match A or C, for instance), and also on exact text matches (such that an M matches with another M but not with A or C). The partial matches similarity is related to the polymorphism p-distance, which assumes that state ambiguity comes from populational diversity (Potts, Hedderson and Grimm, 2014). There is an option for uvaia to mimic other software by excluding partial matches from the comparison ―although we notice that phylogenetic inference methods, specially probabilistic ones, do benefit from partially ambiguous information and thus are also used by default in uvaia.

When we consider indels and Ns, looking at the distance between sequences may not be a good indicator of neighbourhood since the sequences may have few comparable sites (pairwise comparisons exclude sites with a gap or N in one of the sequences, see Figure 1 for an example). The same caveat applies to percentage identity calculation, since we normalise by valid pairwise comparisons. Thus our “neighbourhood” (groups of similar genomic sequences) is defined by the total number of matches, instead of number of mismatches or fraction of matches.

**Figure 1:**
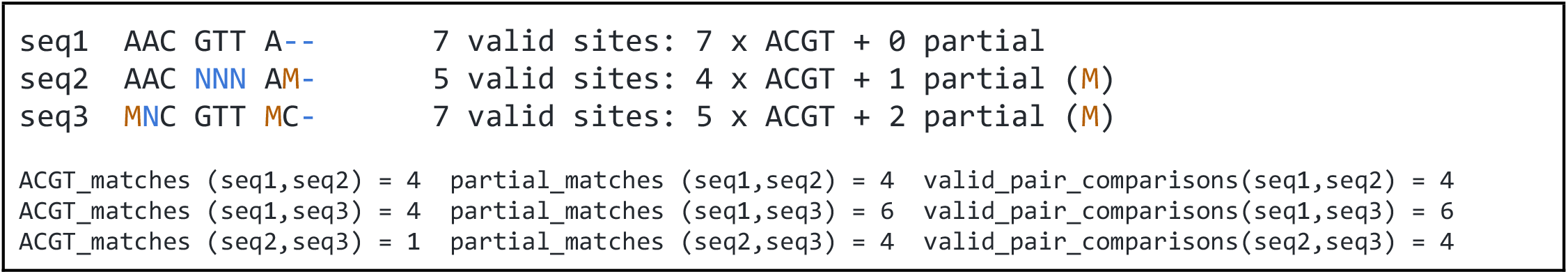
Example of three sequences with no dissimilarities to each other (zero distance) which nonetheless contain differences.

## Methods

Given a set of query sequences, we want to keep from a (potentially very large) reference alignment only the sequences which are close to at least one query. This keeps downstream inferences computationally feasible, and also helps faster inferences to be made, like neighbour-based lineage classification, or geographical analyses (Mashe, Takawira, de Oliveira Martins, *et al*., 2021; Sarwar *et al*., 2021; Eales, de Oliveira Martins, *et al*., 2022). In particular, in uvaia a priority queue is created to store the neighbourhood of each query sample, where at most *k* reference sequences are kept. For each query, a new reference is added to its queue, in order of importance, by its total number of unambiguous matches, of exact text matches, and of partial matches. Ties are further broken by the number of derived unambiguous matches, and ultimately by the number of valid sites of the reference sequence. The reference sequences are thus ranked for each query according to the tie-breaking statistics described above. The number of derived matches is based on the strict consensus between all queries, which is created to speed up calculations and split the sites into constant and polymorphic. The total number of matches is then the sum of ancestral matches (i.e. over sites where all queries have the same state) and derived matches (over sites that may differ between queries). The rationale is to give preference to neighbours closer to the tips and farther from the ancestor of the query sequences.

The same reference can be on the neighbourhood of more than one query sequence, but to speed up computations and to minimise duplicated effort, uvaia can remove identical and redundant query sequences. A sequence is redundant if there is another, more resolved sequence, with all its information but with less ambiguity. For example the 6-mer AACNNN is redundant with respect to AACAAA since the latter is a more resolved version of the former. Any close neighbour to the more resolved sequence will also be a close neighbour to the less resolved one. Notice that on the other hand NACAAA is not a more resolved version of AACNNN since there is information in the latter (the first “A”) not available in the more resolved, former sequence.

At the end, a table with match information between each query and its set of N closest neighbouring reference sequences is output, together with a reduced reference alignment output with all temporary close neighbours. This reduced alignment file can be queried afterwards using the table information, to obtain the neighbour sequence themselves. Uvaia can also simulate other SNP distance algorithms where partially ambiguous sites are not taken into account (considering only A, C, G, and T).

The main program is thus uvaia, which uses the match similarities described above to, given a query data set of aligned sequences, ranks and extracts the closest neighbours in a large reference data set. We also have incorporated an implementation of the wavefront alignment algorithm (Marco-Sola *et al*., 2020) into a reference-based aligner, called uvaialign. This program is also multithreaded, works in batches to avoid using up all available memory up, and can read from and write to compressed format. Uvaialign has been used in several SARS-CoV-2 analyses already (Mashe, Takawira, Gumbo, *et al*., 2021; Sarwar *et al*., 2021; Eales, de Oliveira Martins, *et al*., 2022; Eales, Page, *et al*., 2022; Merhi *et al*., 2022).

## Results

Uvaia can also be used to calculate exhaustively the pairwise similarities between two alignment data sets, and we compared its results to snp-dists (Seemann, 2018). Our test data set is composed of 6000 unique sequences generated at the QIB as part of the COG-UK consortium, spanning the time range 2020-2022, and with the fraction of fully ambiguous sites (i.e. Ns) between 0 and 50%. All sequences were aligned with uvaialign. This data set was split into one set of 1000 samples and one of 5000 samples. Table 1 shows the distribution of PANGO lineages (Rambaut *et al*., 2020) of both sets to give an idea of their diversity. This set was then used to compare uvaia with snp-dists, such that we can calculate the pairwise distance between all sequences in the set.

**Table 1:** PANGO lineages (given by the “scorpio call” column) of the 6000 unique sequences used in this study, divided into two data sets.

|  |  |
| --- | --- |
| 1000 samples data set:<br>341 Omicron (BA.1-like)<br>205 Delta (AY.4-like)<br>196 Probable Omicron (BA.1-like)<br>145 Delta (B.1.617.2-like)<br>55 Other<br>20 Delta (AY.4.2-like)<br>18 Alpha (B.1.1.7-like)<br>10 Omicron (BA.2-like)<br>4 Probable Omicron (BA.2-like)<br>3 Probable Omicron (Unassigned)<br>3 Probable Omicron (BA.3-like) | 5000 samples data set:<br>1725 Omicron (BA.1-like)<br>1096 Delta (AY.4-like)<br>997 Probable Omicron (BA.1-like)<br>719 Delta (B.1.617.2-like)<br>249 Other<br>98 Alpha (B.1.1.7-like)<br>70 Delta (AY.4.2-like)<br>26 Omicron (BA.2-like)<br>10 Probable Omicron (BA.3-like)<br>7 Probable Omicron (BA.2-like)<br>1 Probable Omicron (Unassigned)<br>1 Gamma (P.1-like)<br>1 B.1.617.1-like |

In uvaia we calculate three similarity measures, based on the total number of ACGT matches, total number of text matches (i.e. ACGT plus partially ambiguous treated as characters), and total number of partial matches (where the IUPAC ambiguity code is used to check for compatibility, except for Ns). Thus we can extract the number of mismatches by subtracting the number of pairwise comparisons by these similarity values. In Figure 1 we show cases where they are not equivalent to the SNP distance, since usually these SNP calculations ignore all non-ACGT characters. And therefore even “identical” sequences as reported by e.g. snp-dists may be quite distinct once we look at partially-ambiguous sites. In Figure 2 we have boxplots showing their difference for our smaller data set. Therefore, especially for large databases where the number of sequences without SNPs may be overwhelming, we need higher resolution exclusion criteria for neighbours search.

**Figure 2.**
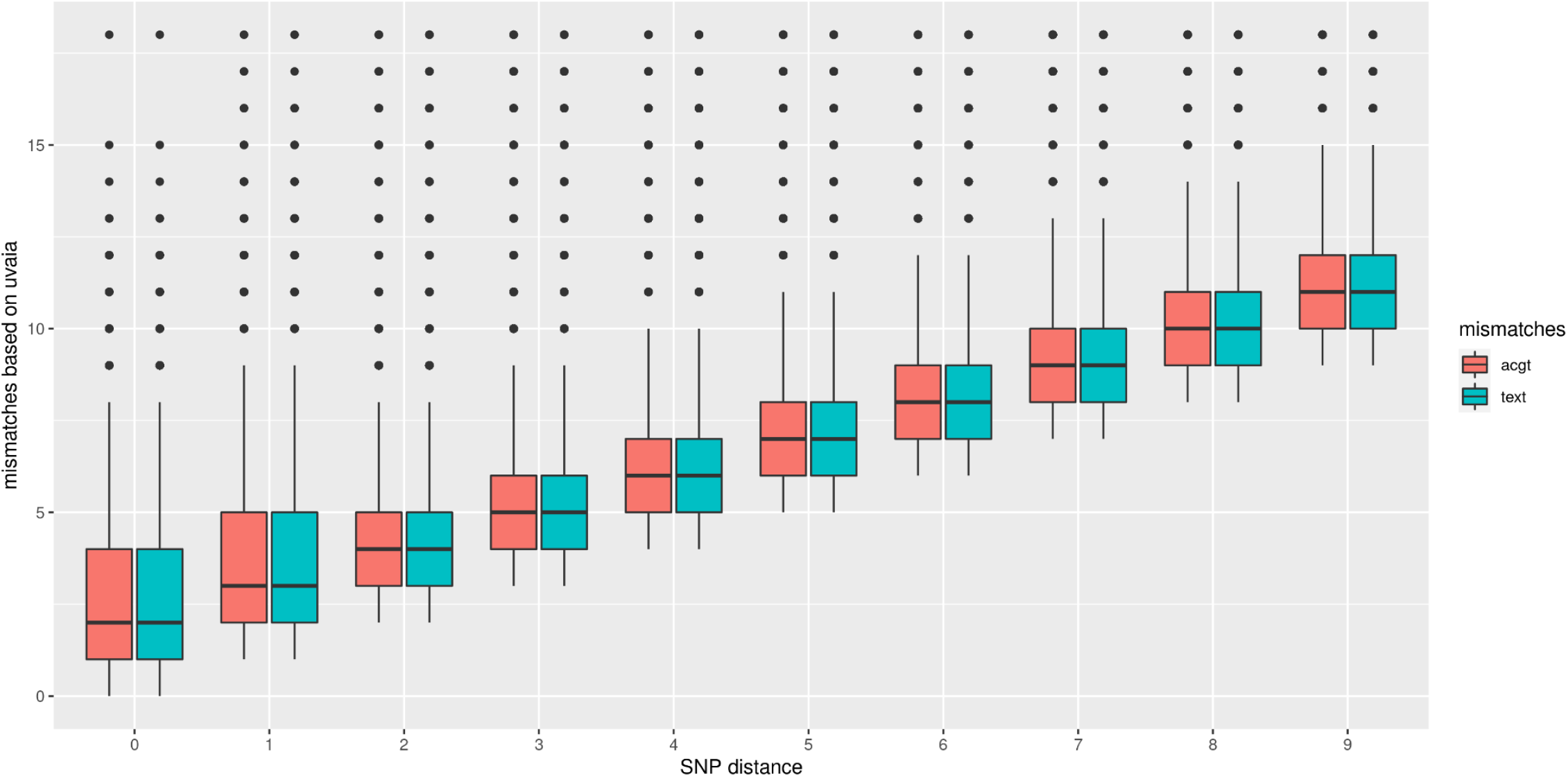
Comparison between the pairwise sequence distance as calculated by snp-dists (x axis) and the number of uvaia mismatches as the difference between the number of valid pairwise comparisons and the number of matches (y axis). The y axis is truncated at the 95% percentile to ease visualisation. We used a set of 1000 sequences, such that 499500 pairwise comparisons are shown.

In Figure 3 we show that by using partial matches we have more similar distance measures. The exceptions are generally cases where there is a comparison between an unambiguous site and an incompatible ambiguous one, as for example between T in one sequence and R in the other (R is compatible with A and G, but not T). In these cases the distance based on partial matches will be higher than the SNP distance (which excludes such sites from the comparison).

**Figure 3.**
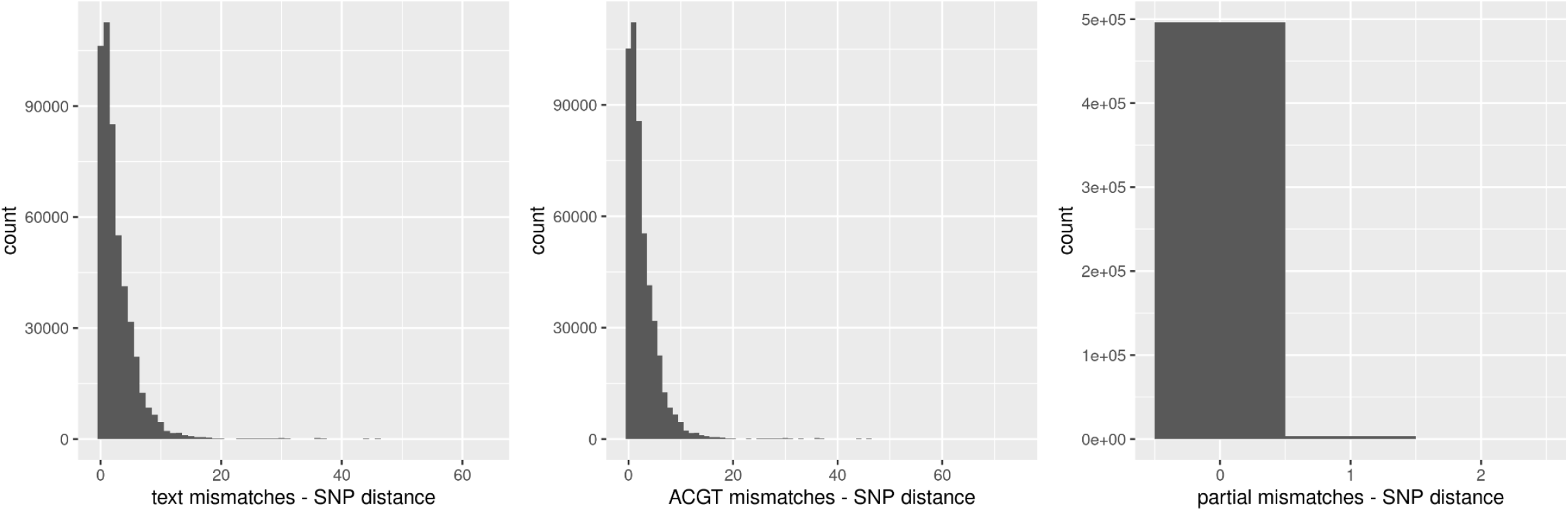
Histogram of the difference in mismatches between uvaia and snp-dists. While their difference is quite evident when we compare ACGT-only or text mismatches, the number of partial mismatches is almost identical to the SNP distance, differing at most by one in approx 3000 pairs, out of the 499500 comparisons).

Uvaia can replicate snp-dists results by excluding partially ambiguous states, i.e. treating them together with Ns and gaps. We then used uvaia to calculate the similarity between these 1000 “query” sequences to 5000 distinct “reference” sequences with similar distribution of lineages and ambiguous sites. To show the effect of uncertainty, we compare the number of neighbours with no SNPs to the reference sequences to their number of (partially) unambiguous sites in Figure 4. We see how less resolved sequences appear to have more “identical” neighbours, i.e. sequences with no SNP differences. Thus, by using the number of matches instead of SNP differences, we can account for this ambiguity.

**Figure 4.**
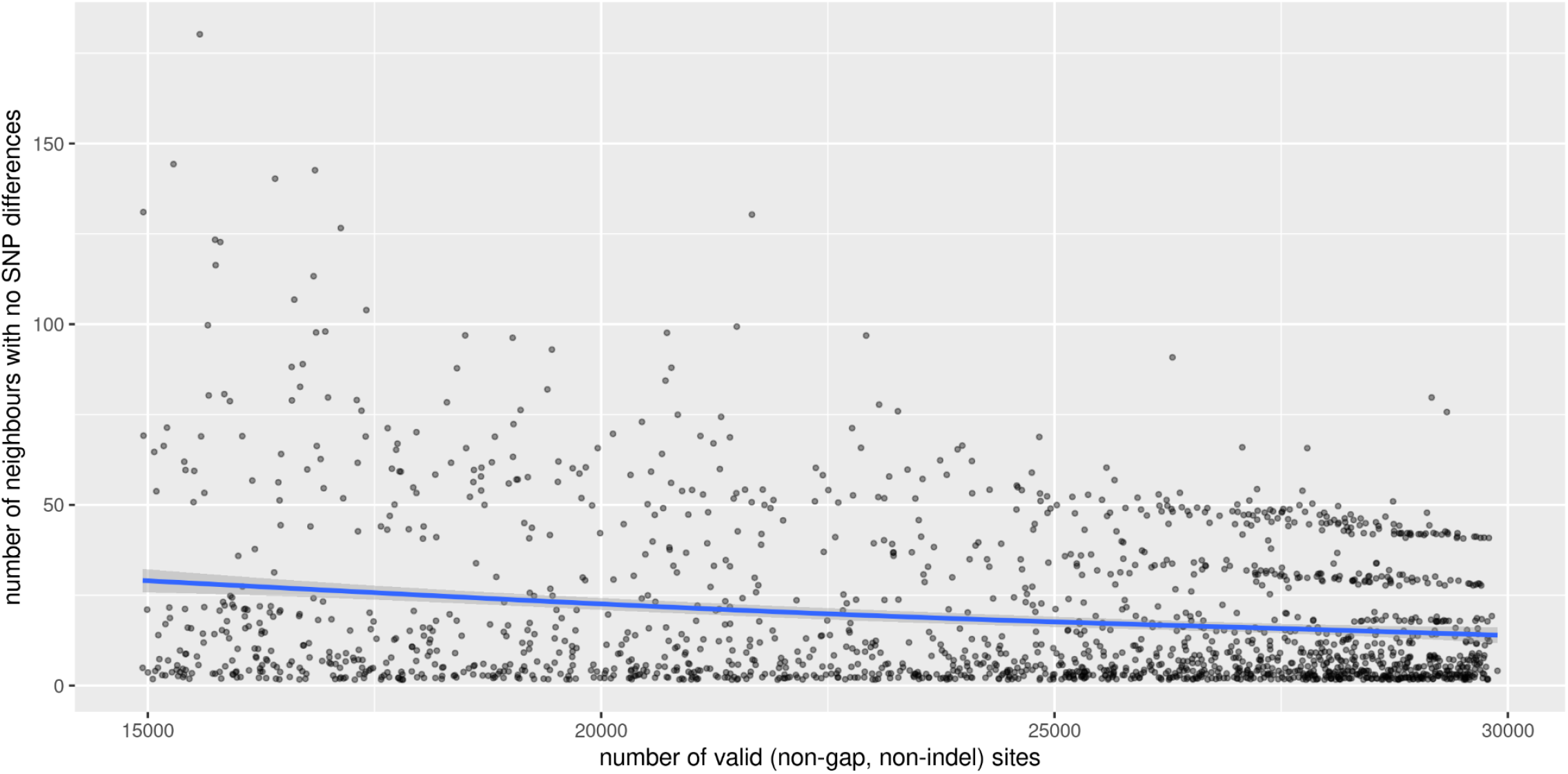
Scatterplot of number of “query” neighbours with no SNPs to “reference” sequences (y axis) against number of partially or completely unambiguous sites in the “reference” sequence (x axis). The blue line represents the regression line smoothed with a generalised additive model (Wood, 2017).

## Discussion

It is more likely to find “identical” neighbours for poorly-resolved sequences, if by identical we mean there are no SNPs or there is 100% identity or similarity (amongst the comparable sites). Therefore we must also account for the number of at least partially unambiguous sites in the comparison, by (1) restricting our comparison to well-resolved sequences (i.e. where the fraction of unambiguous sites is negligible) or by (2) looking at the matches instead of mismatches, focusing on the similarity instead of the distance. Most research in SARS-COV2 has employed strategy (1), by including only sequences with e.g. 90% unambiguous sites. Here we provide a software to solve (2), which can return the reference sequences with more matches to a query sequence.

This may affect not only the phylogenetic inference ―since likelihood and Bayesian methods can use the ambiguity information― but also in other epidemiological analyses where finding the closest sequences may be challenging. Uvaia has been used successfully in both cases, by improving contact tracing and travel history inferences, and overall evolutionary analyses (Mashe, Takawira, de Oliveira Martins, *et al*., 2021; Mashe, Takawira, Gumbo, *et al*., 2021; Sarwar *et al*., 2021; Aggarwal *et al*., 2022; Eales, de Oliveira Martins, *et al*., 2022; Merhi *et al*., 2022), and here we present further details into the algorithm.

Some pipelines might inadvertently replace ambiguous sites by the reference allele, mistakenly assuming that low coverage sites are evidence for no change (the ancestral state). This imputation strategy may inadvertently lead to misleading inferences since, as indicated by Figure 4, it can negatively affect similarity and distance estimates. Furthermore it can mask the effects of recombination, intra-population diversity, homoplasies, and can mislead phylogenetic analyses (Baker *et al*., 2021).

## Conclusion

We show how a SNP distance-based neighbour sequence search may have low resolution to find the most similar sequences in large databases. Notice that the main differences between uvaia and other algorithms is the inclusion of partially ambiguous sites (with the possibility of incorporating or not their compatibility information), and the number of matching sites as optimality criterion. It is known that such partial matches can affect phylogenetic analyses (Potts, Hedderson and Grimm, 2014), and a so-called “Intra-Individual Site Polymorphism” is available in the R library phangorn (Schliep, 2011).

We note that working with compressed files has a computational cost, even when using a multithreaded algorithm for compression; decompression is always single threaded, but faster than compression. However we have an overall time benefit if we always store the files compressed, due to their large sizes. The advantages of uvaia are best seen in a restricted budget environment, with finite disk and memory resources, but fully using multicore architectures available even in lower end laptops. And as we are already seeing for SARS-CoV-2, in preparation for future pandemics, new software like uvaia will be the default, scalable not for thousands but for millions of sequences.

## Ethics

The COVID-19 Genomics UK Consortium has been given approval by Public Health England’s Research Ethics and Governance Group (PHE R&D Ref: NR0195).

## Acknowledgements

The authors would like to thank the Norfolk and Norwich University Hospitals NHS Foundation trust for providing samples for sequencing, and the Bioinformatics and Sequencing teams at the Quadram for generating all data analysed in the present study.

## Author contributions

First draft: LdOM Funding acquisition: AJP

Leadership and supervision: AEM, AJP Sequencing and analysis: LdOM Visualisation: LdOM

## Data availability

All sequence read data and consensus genome sequences are available without restriction from the European Nucleotide Archive at https://www.ebi.ac.uk/ena/browser/view/PRJEB37886, and consensus genome sequences (above 90% genome coverage) are available from the Global initiative on sharing all influenza data at https://www.gisaid.org, with accession numbers for each sample listed in Supplementary Table 1. All data are also available in https://github.com/quadram-institute-bioscience/uvaia.

## Availability

Uvaia and uvaialign are available through github on https://github.com/quadram-institute-bioscience/uvaia, and also on conda and docker. The source code is distributed under a GPL3 licence.

